# Microbial neurosteroid biosynthesis: *Holdemania filiformis* generates isopregnanolone that reaches the brain via gut-to-brain transport

**DOI:** 10.1101/2024.12.12.628284

**Authors:** Guo-Jie Brandon-Mong, Chia-Hung Chou, Tien-Yu Wu, Ronnie G Gicana, Chien-Tao Huang, Guan-Lun Wu, Ya-Che Li, Yi-An Tu, Yu-Yuan Chang, Yi-Li Lai, Po-Ting Li, Hui-Yu Chen, Tsun-Hsien Hsiao, Mei-Jou Chen, Yin-Ru Chiang

## Abstract

5α-neurosteroids such as allopregnanolone and isopregnanolone play critical roles in neurological health and mood regulation, yet current therapeutic production faces significant limitations. We demonstrate that specific gut microbes represent a previously unrecognized source of bioavailable 5α-neurosteroids that reach the central nervous system via the gut-brain axis. Through integrated metabolomic and genomic analyses of progesterone-amended fecal cultures, we identified *Holdemania* as a major producer of isopregnanolone via microbial steroid 5α-reductase (BaiJ type 2) and 3β-hydroxysteroid dehydrogenase/reductase. Phylogenetic analysis revealed that BaiJ-like sequences cluster predominantly within Firmicutes, with *Holdemania* species forming a distinct clade. In female C57BL/6 mice administered progesterone and *H. filiformis*, 5α-neurosteroids including isopregnanolone predominated in gut tissues while allopregnanolone was the major hepatic neurosteroid. Critically, using stable isotope tracing with [3,4-¹³C₂]progesterone, we detected ¹³C-labeled isopregnanolone in brain tissue, providing direct evidence for gut-to-brain transport of microbiota-derived neurosteroids. High-fat diet significantly enhanced brain 5α-neurosteroid accumulation. Global meta-analysis reveals reduced *Holdemania* abundance in PCOS patients (n = 346) compared to healthy women (n = 321). These findings identify gut microbiota as pharmacologically relevant neurosteroid producers and position *H. filiformis* as a promising probiotic candidate for enhancing endogenous neurosteroid production to treat mood disorders and other neuropsychiatric conditions.

**Highlights:**

- *Holdemania* was identified as a major producer of 5α-neurosteroids, particularly isopregnanolone, in the intestinal tract
- *Holdemania* 5α-reductase (BaiJ type 2) belongs to a distinct phylogenetic clade compared to characterized *Clostridium* BaiJ (type 1)
- 5α-neurosteroids occurred predominantly in the cecum of female mice administered *H. filiformis*, progesterone, and high-fat diet
- ¹³C-labeled 5α-neurosteroids were detected in brain tissue of female mice orally administered [3,4-¹³C₂]progesterone and *H. filiformis*, demonstrating gut-to-brain transport
- Gut microbes such as *H. filiformis* represent promising probiotic candidates for enhancing 5α-neurosteroid production and circulation via the gut-brain axis

**Graphical Abstract:** 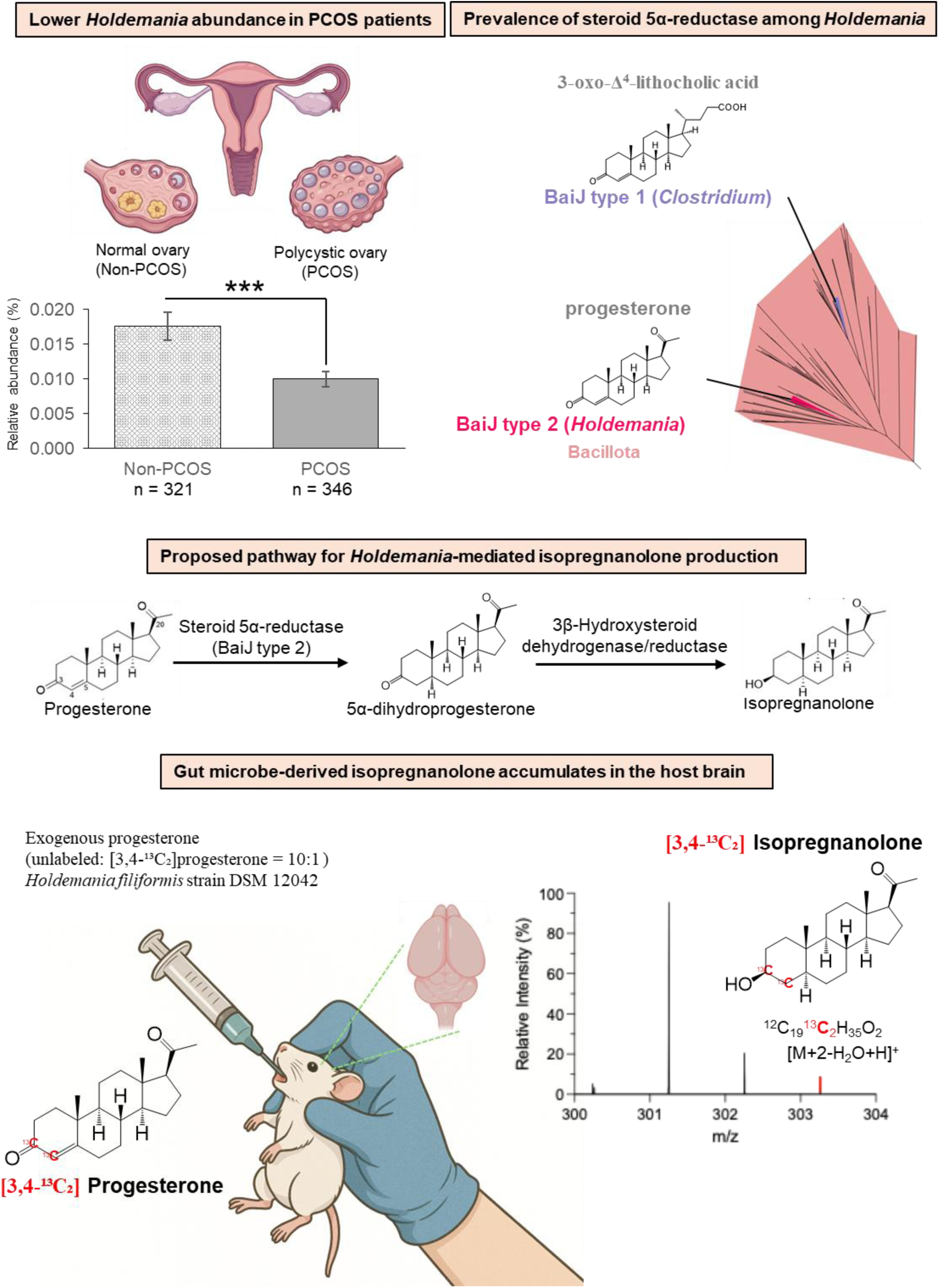

## INTRODUCTION

Neurosteroids (with 19 to 21 carbons) represent a class of potent neuromodulatory compounds that profoundly influence central nervous system function, including neurotransmission, neuroplasticity, and behavior.^1,2^ Progesterone-derived C_21_ neurosteroids, such as allopregnanolone, isopregnanolone, pregnanolone, and epipregnanolone (see **Figure 1** for individual structures), have emerged as critical molecules in neuroscience and medicine due to their profound impacts on human health through modulation of γ-aminobutyric acid type A (GABA_A_) receptors, influencing neuronal excitability and various neurophysiological processes.^3–5^ In humans, the conversion of progesterone to allopregnanolone involves sequential enzymatic reactions catalyzed by human steroid 5α-reductases (SRD5A1 and SRD5A2) and 3α-hydroxysteroid dehydrogenase/reductase, predominantly expressed in the liver.^6,7^ However, whether hepatically produced allopregnanolone efficiently crosses the blood-brain barrier remains unclear, which may restrict its neuromodulatory effects in the central nervous system.^8^

**Figure 1.**
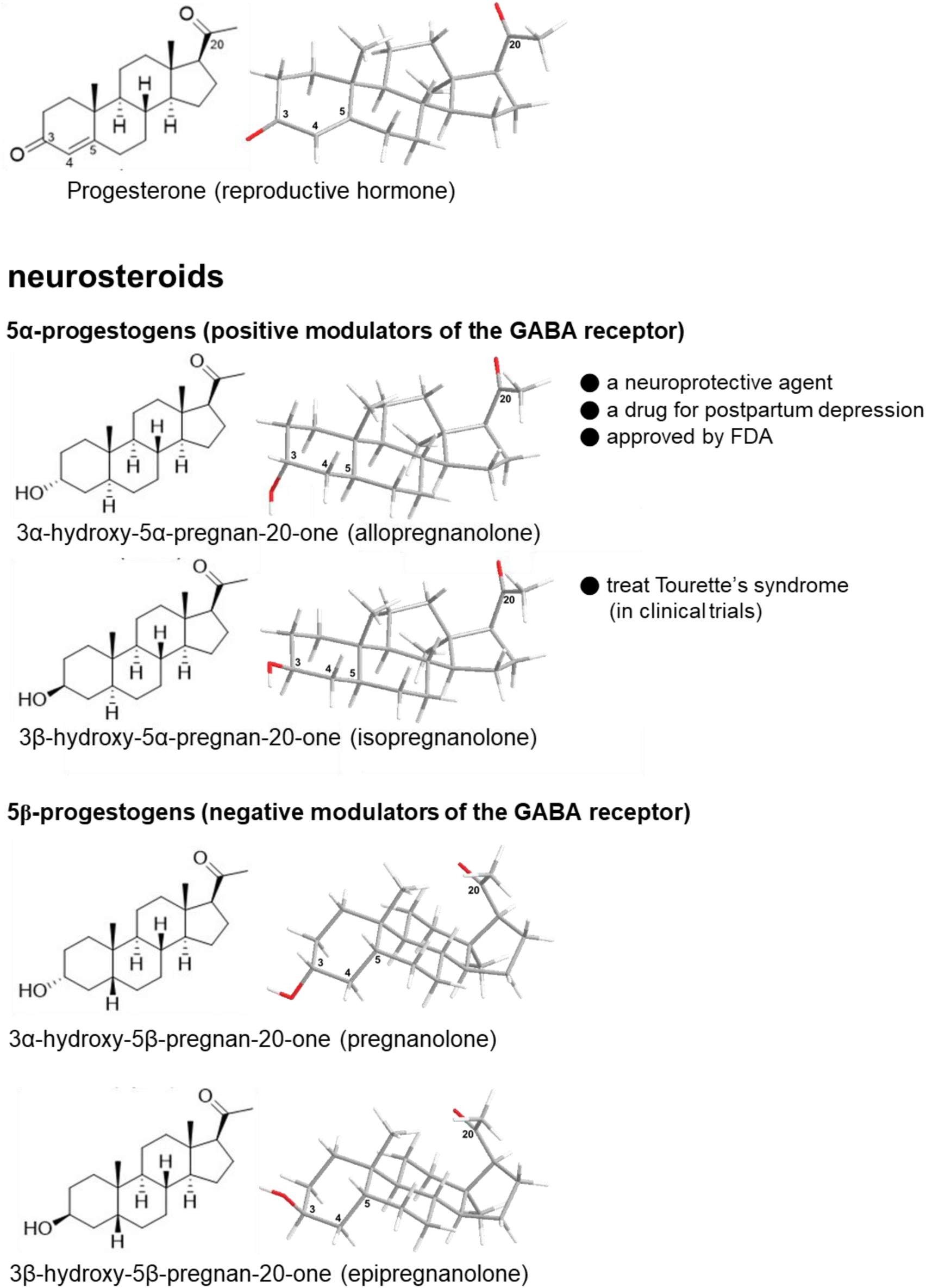
Two- and three-dimensional structural representations of progesterone and its neurosteroid derivatives. The structures include progesterone, allopregnanolone (5α3α), isopregnanolone (5α3β), pregnanolone (5β3α), and epipregnanolone (5β3β). The 5α-progestogens (allopregnanolone and isopregnanolone) and 5β-progestogens (pregnanolone and epipregnanolone) are distinguished by their A/B-ring conformations. Key carbon atoms are numbered on the progesterone structure and its derivatives according to the standard steroid numbering system.

The clinical significance of neurosteroids has been underscored by recent therapeutic breakthroughs. The global neurosteroid therapeutics market is experiencing unprecedented growth, driven by increasing recognition of these molecules’ therapeutic potential and the FDA approvals of allopregnanolone (Zulresso™) for postpartum depression in 2019 and zuranolone (an allopregnanolone derivative) as the first oral antidepressant in 2023.^9,10^ However, current manufacturing approaches face significant challenges that limit market accessibility and therapeutic development. Conventional neurosteroid production relies on either isolating steroids from animal tissues or total chemical synthesis that lacks stereoselectivity, resulting in mixed stereoisomer populations requiring laborious and costly purification procedures.^11–14^

Recent discoveries have fundamentally challenged the traditional view that neurosteroid biosynthesis is exclusively a host-driven process. Specific gut microorganisms possess sophisticated enzymatic machinery capable of steroid hormone biotransformation. For instance, *Ruminococcus gnavus* and *Bacteroides acidifaciens* can convert pregnenolone and hydroxypregnenolone into dehydroepiandrosterone and testosterone, respectively,^15^ while bacterial genera *Gordonibacter* and *Eggerthella* demonstrate the capacity to synthesize neurosteroids from gut-derived glucocorticoids.^16^ Our previous research further demonstrated that *Clostridium innocuum* metabolizes progesterone into epipregnanolone via sequential 5β- and 3β-reduction.^17^ Collectively, these findings establish the gut microbiota as a previously underappreciated but potentially significant source of circulating neurosteroids, with important implications for understanding hormone metabolism and therapeutic outcomes.

Elucidating bacterial contributions to host neurosteroid profiles presents significant methodological challenges. The relevant bacterial species comprise less than 0.1% of the gut microbiota,^18,19^ and the complex interplay between human and microbial enzymatic pathways in neurosteroid biosynthesis remains poorly understood. Additionally, the transport efficiency and mechanisms by which gut-derived neurosteroids reach the brain have not been established. To address these knowledge gaps, we developed an integrated multi-omics approach to investigate gut microbiota-mediated neurosteroid production. Human fecal cultures were supplemented with progesterone to selectively enrich progesterone-metabolizing gut microbes, and the resulting neurosteroid profiles were characterized using Ultra-Performance Liquid Chromatography coupled with High-Resolution Mass Spectrometry (UPLC–HRMS). Functional genomics approaches were employed to assess the prevalence of progesterone-to-neurosteroid transformation genes among gut microbiota.

Our results demonstrate that specific gut microbes, including *Holdemania* species, serve as alternative sources of 5α-neurosteroids through novel biosynthetic mechanisms. Critically, we provide direct evidence for gut-to-brain transport of microbiota-derived neurosteroids: ¹³C-labeled neurosteroids were detected in brain tissue of female mice following oral administration of [3,4-¹³C₂]progesterone and *H. filiformis*. Notably, the neurosteroid profiles in mouse brain tissue showed greater similarity to gut tissue patterns than to hepatic profiles, supporting preferential gut-brain transport pathways. These findings identify gut microbes such as *H. filiformis* as promising probiotic candidates for therapeutic enhancement of 5α-neurosteroid production and bioavailability through targeted modulation of the gut-brain axis.

## RESULTS

### Characterization of key metabolites produced by gut microbiota during anaerobic progesterone metabolism

To investigate progesterone-to-neurosteroid transformation, we incubated fecal samples from 14 women (**Table S1**) with 1 mM progesterone under anaerobic conditions using two distinct media: Brain Heart Infusion (BHI; a nutrient-rich medium) and DCB-1 (a chemically defined, nutrient-poor medium).^20^ Residual progesterone and its metabolites were analyzed using UPLC–atmosphere pressure chemical ionization (APCI)–HRMS (**Figure 2**). In the nutrient-rich BHI medium, gut microbiota rapidly metabolized over 90% of progesterone within 3 days. This biotransformation generated multiple neurosteroids: allopregnanolone (0.05 mM), epipregnanolone (0.06 mM), isopregnanolone (0.07 mM), and pregnanolone (0.65 mM) (**Figure 2A**, left panel). Conversely, in the nutrient-poor DCB-1 medium, approximately 0.7 mM (30%) of progesterone remained unchanged, with pregnanolone as the predominant metabolite (**Figure 2A**, right panel).

**Figure 2.**
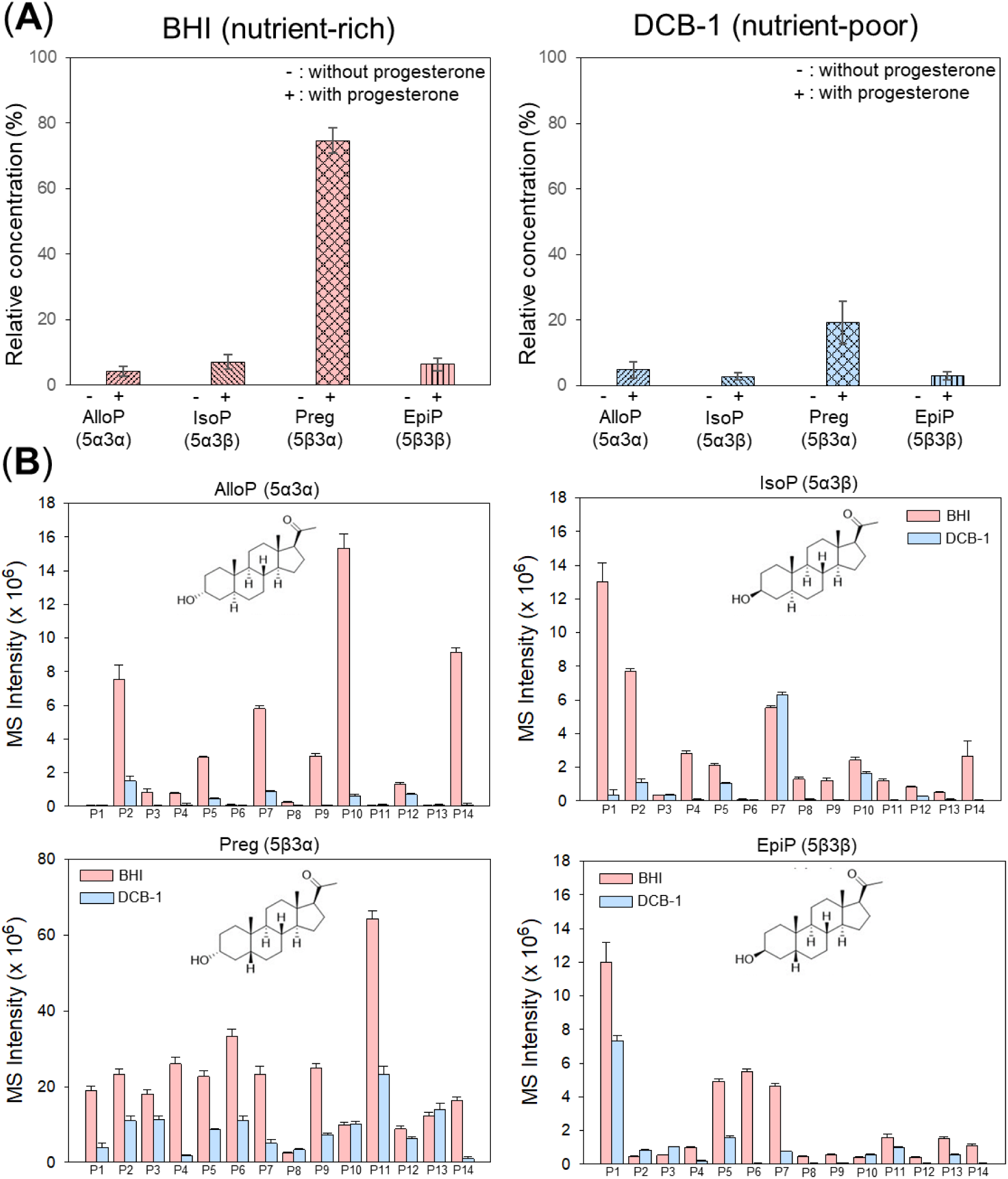
Metabolomic analysis of fecal cultures from progesterone-treated women under anaerobic conditions. Fecal samples were collected from 14 females receiving progesterone treatment (P1-P14) and cultured in nutrient-rich (BHI) or nutrient-poor (DCB-1) media supplemented with exogenous progesterone (1 mM). **(A)** Quantification of major C_21_ neurosteroids in BHI (left) and DCB-1 (right) cultures supplemented with progesterone. Values represent mean ± SEM (n = 14). **(B)** Distribution of major microbial metabolites across individual samples in progesterone-supplemented cultures. Abbreviations: IsoP, isopregnanolone; AlloP, allopregnanolone; Preg, pregnanolone; EpiP, epipregnanolone.

Individual-specific analysis revealed distinct neurosteroid production patterns (**Figure 2B**). Most fecal samples demonstrated enhanced neurosteroid production in BHI compared to DCB-1 conditions, though notable inter-individual variations emerged. Female 11’s microbiota exclusively produced pregnanolone, while Female 1’s microbiota generated epipregnanolone, isopregnanolone, and pregnanolone but not allopregnanolone. Females 10 and 14’s microbiota predominantly synthesized allopregnanolone and pregnanolone. Despite this heterogeneity, pregnanolone consistently emerged as the dominant metabolite across all samples.

These findings demonstrate that gut microbiota primarily converts progesterone to 5β-neurosteroids, particularly pregnanolone—a pattern observed consistently across our female cohort. Notably, our metabolite profile analysis reveals the presence of gut microbes capable of anaerobic transformation of progesterone into 5α-neurosteroids, including allopregnanolone and isopregnanolone—compounds with significant therapeutic potential.^2,8–10^

### Progesterone-induced changes in gut microbial composition

To identify the major bacterial species involved in anaerobic progesterone metabolism, we analyzed bacterial community structures in progesterone-amended fecal cultures through DNA extraction, 16S rRNA gene amplification, and PacBio sequencing. Non-metric multidimensional scaling (NMDS) analysis based on Bray-Curtis dissimilarity revealed distinct changes in the progesterone-treated gut microbiota. In the nutrient-rich BHI broth, baseline control samples clustered at the top right of the coordinate, while samples from the progesterone-amended cultures stage clustered at the bottom (**Figure 3A**, left panel). Permutational multivariate analysis of variance (PERMANOVA) confirmed significant differences (global *R*^2^ = 0.34; *p* = 0.001) between baseline control and progesterone-enriched gut microbiota (**Figure 3A**, left panel). In BHI cultures incubated with progesterone, we observed significant increases in Bacillota (synonym Firmicutes; 69.1%; *p* < 0.05) (**Figure 3B**, left panel) and Pseudomonadota (synonym Proteobacteria; 11.3%; *p* < 0.01) (**Figures 3B** and **S1**).

**Figure 3.**
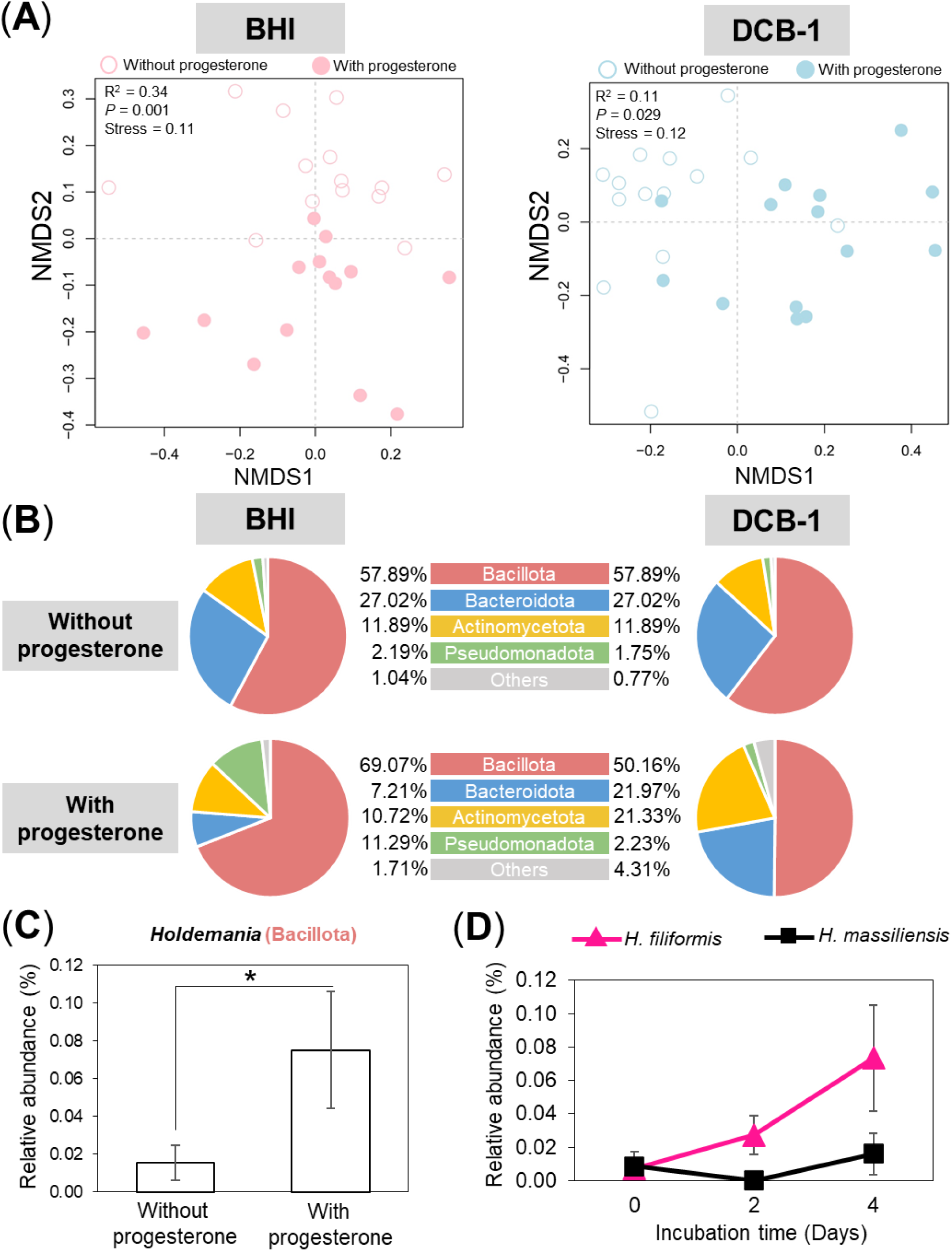
Progesterone treatment alters fecal bacterial community structure. In the progesterone-enriched cultures, fecal samples from 14 female patients were cultured anaerobically with 1 mM progesterone in nutrient-rich BHI medium (left panels) or nutrient-poor DCB-1 medium (right panels), and analyzed by PacBio sequencing. (**A**) NMDS ordination demonstrates significant separation of bacterial communities between progesterone-enriched and baseline samples. (**B**) Certain bacterial phyla were significantly enriched following progesterone treatment. see **Figure S1** for statistical analysis. **(C)** *Holdemania* was significantly enriched following progesterone treatment (see **Figure S2** for the top 30 enriched genera). **(D)** Temporal dynamics of *Holdemania* species abundance, particularly *H. filiformis*, in progesterone-treated fecal cultures. Data represent mean ± SEM (n = 14). Statistical significance was determined using the Wilcoxon rank-sum test at the phylum level and DESeq2 at the genus level; *p < 0.05.

In the nutrient-poor DCB-1 broth, control samples clustered at the top left, while progesterone-supplemented cultures clustered on the right (**Figure 3A**, right panel). PERMANOVA showed a modest but significant difference (global *R*^2^ = 0.11; *p* = 0.029) between DCB-1 control and progesterone-enriched communities (**Figure 3A**, right panel). After anaerobic incubation in this nutrient-poor medium, three bacterial phyla showed significant increases: Actinomycetota (synonym Actinobacteria; *p* < 0.05), Thermodesulfobacteriota (synonym Thermodesulfobacteria; *p* < 0.01), and Synergistota (synonym Synergistetes; *p* < 0.05) (**Figures 3B and S1**).

At the genus level, the relative abundance of *Holdemania*, a member of Bacillota, increased notably following anaerobic incubation with progesterone (**Figure 3C**; see **Figure S2** for other enriched genera). *H. filiformis* abundance increased markedly during anaerobic growth with progesterone. A similar trend was observed for *H. massiliensis*, although to a lesser extent (**Figure 3D**).

### Progesterone-to-5α-neurosteroid transformation genes across gut bacterial genomes

Metabolomic analysis of progesterone-amended fecal cultures demonstrated that gut microbiota can produce the complete spectrum of C_21_-neurosteroids. Bacterial community structure analysis revealed that members of Bacillota represent the predominant gut microbes responsible for anaerobic neurosteroid biosynthesis. Given that 5α-neurosteroids, including allopregnanolone and isopregnanolone, function as positive allosteric modulators of the GABA_A_ receptor and therefore possess significant therapeutic potential for mood disorders,^2–5^ we sought to identify gut microbes—particularly Bacillota species—capable of catalyzing progesterone-to-5α-neurosteroid conversion.

The steroid 5α-reductase BaiJ, involved in microbial steroid metabolism, has been biochemically characterized in *Clostridium scindens*, a prevalent gut Bacillota species.^21^ While the substrate specificity of BaiJ for progestogens remains undefined, gut microbes harboring BaiJ homologs represent promising candidates for 5α-neurosteroid biosynthesis. Using *C. scindens* BaiJ as a query sequence, we identified 17 bacterial species containing BaiJ homologs with >45% sequence identity (**Figure 4**). This scarcity of putative 5α-neurosteroid producers aligns with our metabolomic findings, which identified pregnanolone (5β-neurosteroid) as the predominant microbial metabolite derived from progesterone. Currently, only three *Holdemania* genomes are available in the NCBI database. All three species harbor *baiJ* homologs, suggesting that this genus may play an important role in gut 5α-neurosteroid production. Additionally, all three species encode homologs of 3β-hydroxysteroid dehydrogenase/reductase (3β-HSDR) with >80% sequence identity, indicating that isopregnanolone is likely their primary neurosteroid metabolite. **Phylogenetic distribution of BaiJ-like sequences**

**Figure 4.**
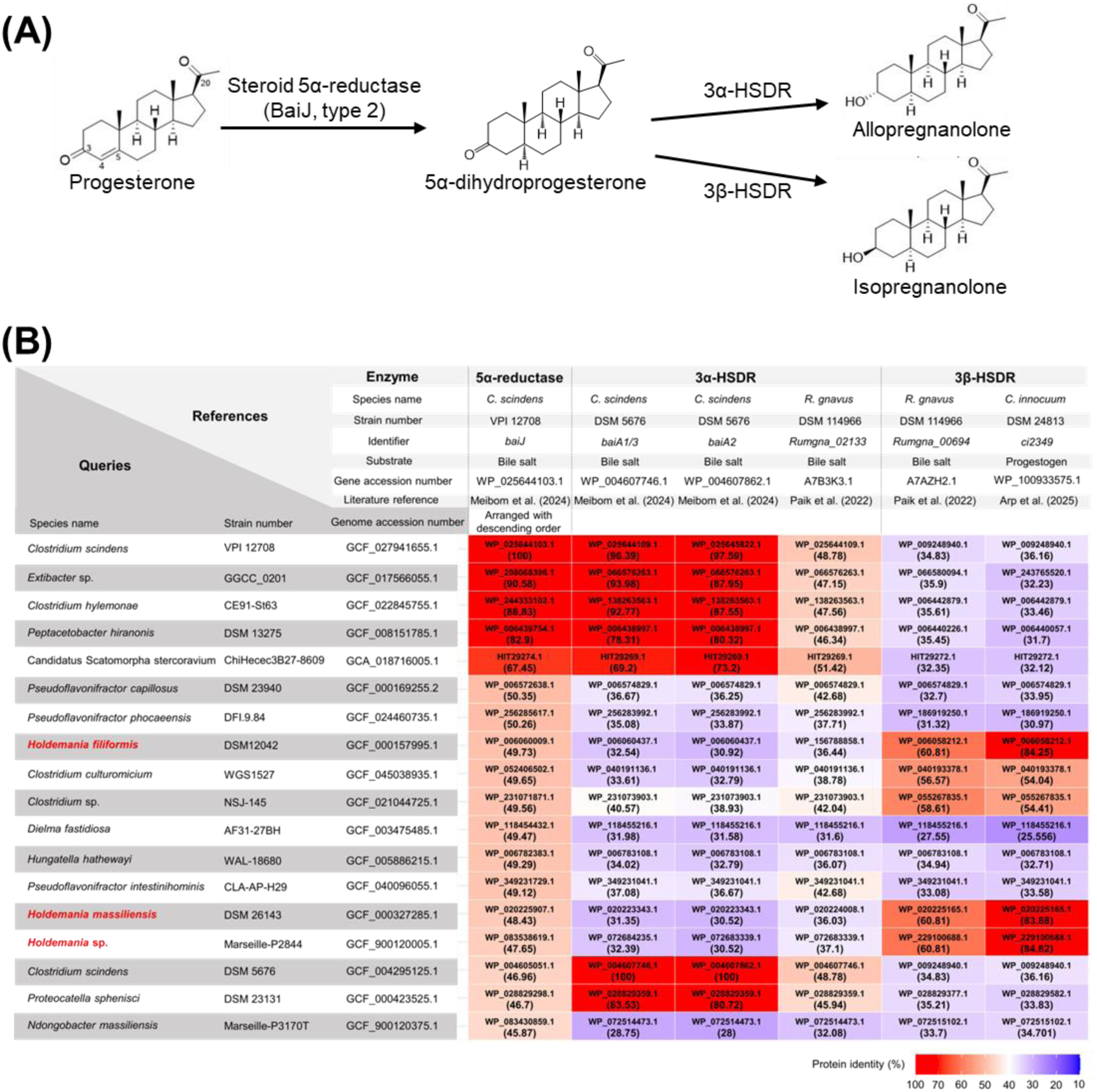
Distribution of steroid 5α-reductase (BaiJ) and hydroxysteroid dehydrogenase/reductase (HSDR) genes across bacterial taxa. (**A**) Microbial enzymatic pathway for biotransformation of progesterone into 5α-neurosteroids. (**B**) Heatmap depicting the distribution of genes encoding 5α-neurosteroid biosynthetic enzymes across bacterial species. BaiJ (type 2) and 3β-HSDR genes are highly prevalent in *Holdemania* species. The *baiJ* gene (type 1) from *Clostridium scindens* VPI 12708, which catalyzes 5α-reduction of 3-oxo-Δ^4^-lithocholic acid, was used as the query sequence for identifying bacterial steroid 5α-reductase genes.

Phylogenetic analysis of BaiJ-like sequences across bacterial phyla revealed distinct evolutionary relationships organized into three major clades (**Figure 5**). Sequences from Actinomycetota (yellow), Bacillota (pink), Bacteroidota (blue), and Pseudomonadota (green) displayed phylum-specific clustering patterns, though some clades contained representatives from multiple phyla. The BaiJ type 1 clade comprised the reference *C. scindens* BaiJ protein along with closely related Clostridia sequences exhibiting 80-90% amino acid identity, including *Clostridium hylemonae* (WP_244333102.1), *Extibacter* sp. GGCC 0201 (WP_208068396.1), and *Peptacetobacter hiranonis* (WP_006439754.1). A second Bacillota-specific cluster, designated BaiJ type 2, contained *Holdemanella filiformis* (WP_006060009.1), *H. massiliensis* (WP_020225907.1), and *Holdemanella* sp. Marseille-P2844 (WP_083538619.1). Node A encompassed additional Bacillota sequences sharing >40% identity with *C. scindens* BaiJ (**Figure 5**).

**Figure 5.**
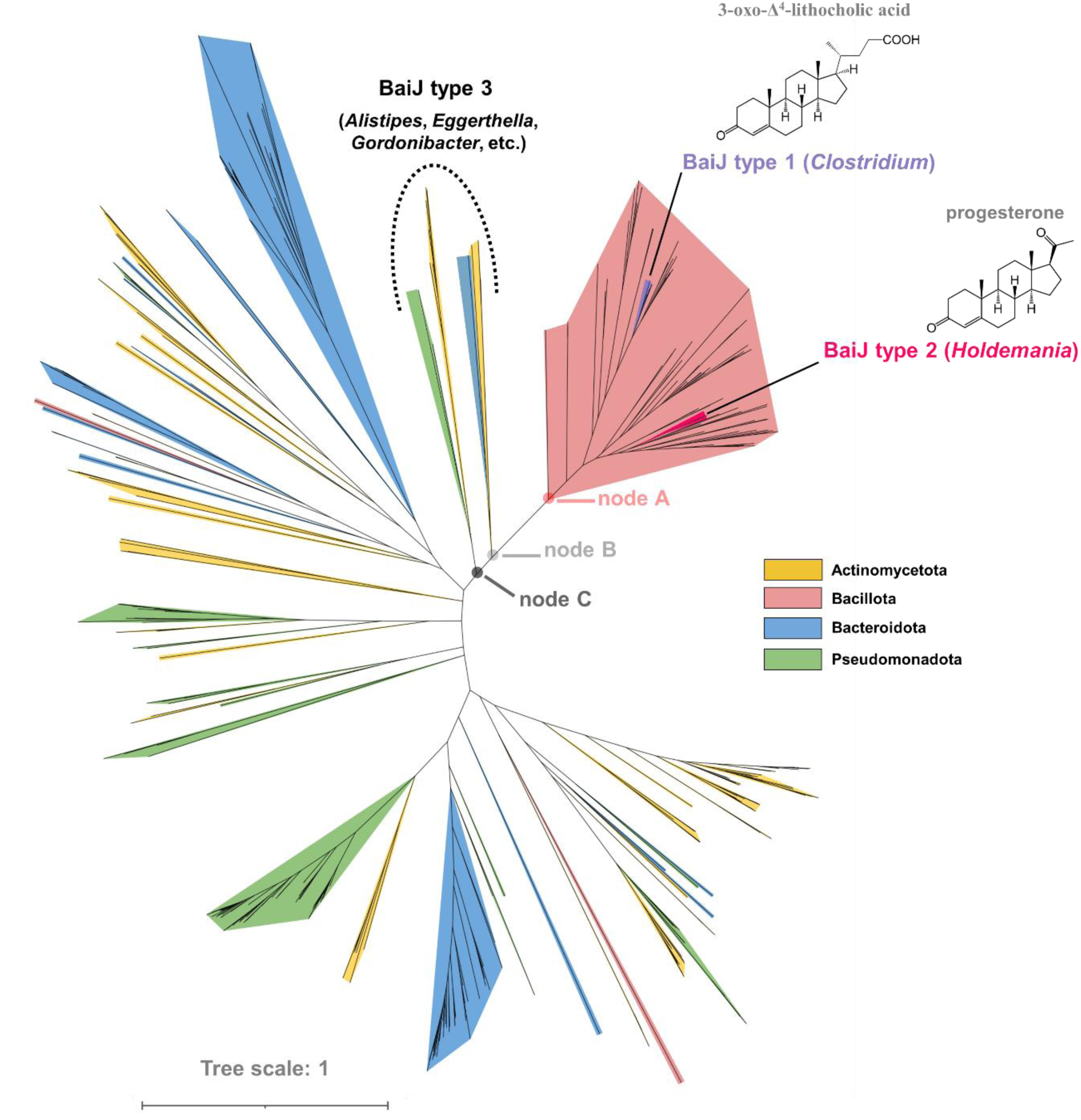
Phylogenetic tree of BaiJ protein sequences across major gut bacterial phyla. Maximum likelihood phylogenetic tree constructed from 390 BaiJ-like protein sequences identified through BLAST searches using *C. scindens* BaiJ as the query sequence. Sequences are colored by phylum: Actinomycetota (yellow), Bacillota (pink), Bacteroidota (blue), and Pseudomonadota (green). The tree reveals three major clades: Type 1 contains the *C. scindens* reference sequence and closely related Clostridia homologs (80–90% amino acid identity); Type 2 includes *Holdemania* species with moderate similarity (48–50% identity; Node A representing sequences with >40% identity); and Type 3 (Nodes B and C) comprises distantly related homologs from Actinomycetota, Bacteroidota, and Pseudomonadota (28–35% identity to *C. scindens* BaiJ).

More divergent BaiJ homologs were identified within the BaiJ type 3 clade, subdivided into nodes B and C. These sequences, derived from Actinomycetota, Bacteroidota, and Pseudomonadota, shared only 28-35% amino acid identity with the *C. scindens* reference sequence (**Figure 5**). Node B included Actinomycetota representatives *Adlercreutzia equolifaciens* (WP_302404602.1), *Gordonibacter* sp. (WP_332711327.1), and *Eggerthella guodeyinii* (WP_195762818.1), alongside *Alistipes* sp. (MBQ3203991.1) from Bacteroidota. Node C contained Pseudomonadota sequences from *Mesosutterella faecium* (WP_243377545.1) and *Janthinobacterium* sp. (MEG1053310.1), together with multiple Actinomycetota species: *Adlercreutzia equolifaciens* (WP_275206498.1), *A. quisgranensis* (WP_160212916.1), *Denitrobacterium detoxificans* (WP_303727570.1), *Eggerthella lenta* (WP_195666307.1), *Curtanaerobium respiraculi* (WP_283170712.1), and Coriobacteriaceae bacterium (MDD5894368.1) (**Figure 5**).

### Confirmation of progesterone-to-isopregnanolone transformation capability of *Holdemania* species

To validate progesterone-to-isopregnanolone conversion capacity, we cultured two *Holdemanella* species (*H. filiformis* strain DSM 12042 and *H. massiliensis* strain DSM 26143, both type strains with sequenced genomes)^22,23^ with 1 mM progesterone under strictly anaerobic conditions. Both strains produced isopregnanolone robustly during anaerobic growth in progesterone-supplemented Peptone Yeast Glucose (PYG) broth (**Figure 6A**), confirming predictions from our functional genomic analysis (**Figures 4** and **5**). Given the higher prevalence of *H. filiformis* in the human gut microbiota,^22,24^ we selected strain DSM 12042 as our model organism for subsequent physiological and metabolic characterization.

**Figure 6.**
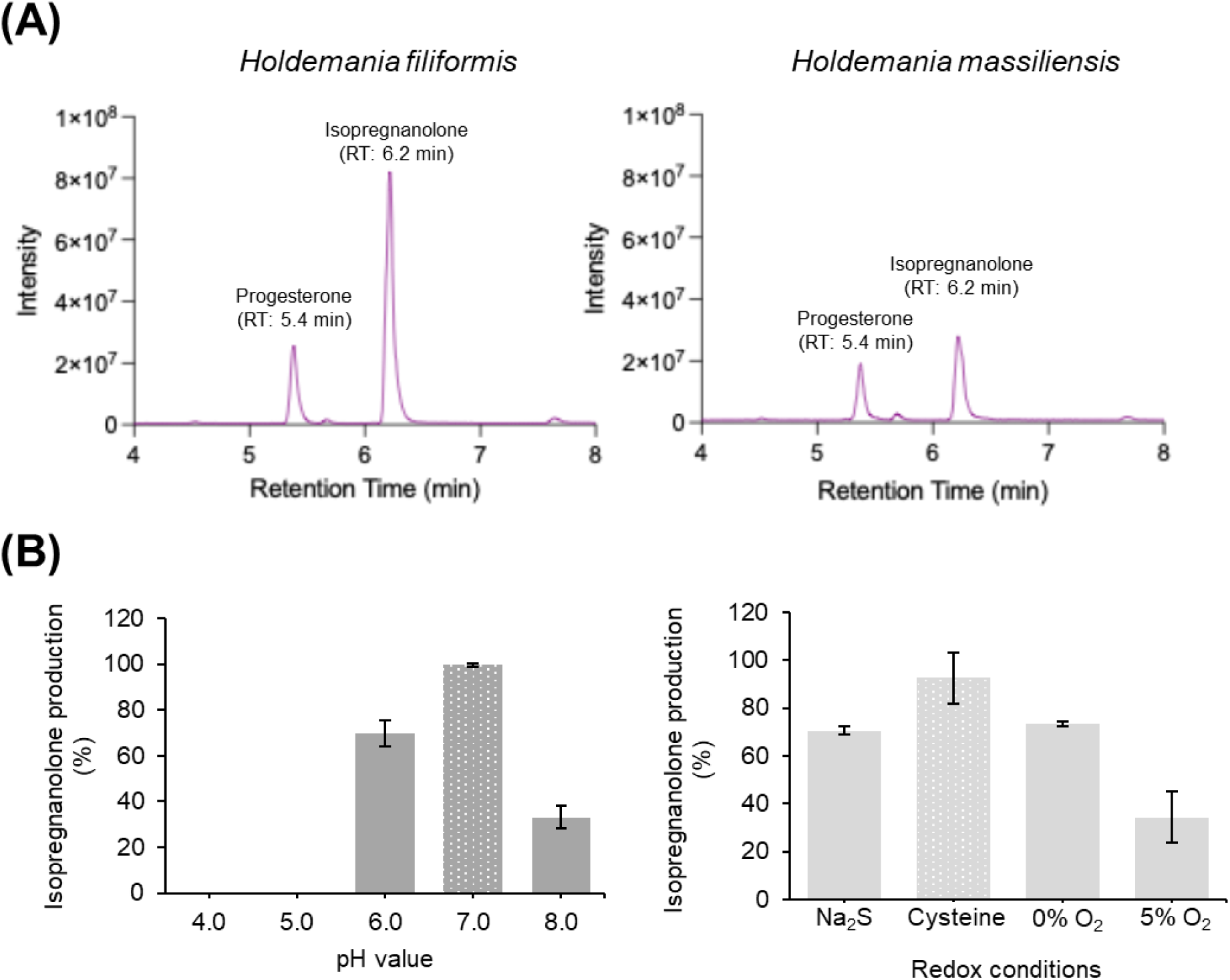
Functional and physiological characterization of isopregnanolone-producing *H. filiformis* strain DSM 12042. (**A**) Both *Holdemania* species exhibit comparable isopregnanolone production. (**B**) Optimal conditions for *H. filiformis*-mediated conversion of progesterone to isopregnanolone. In pH optimization experiments (left panel), 1 mM cysteine was included as a mild reducing agent. In experiments testing oxygen availability and redox conditions (right panel), pH was maintained at 7.0 across all conditions. Data are presented as mean ± SEM from three independent experiments and normalized to 100% relative to strain DSM 12042 cultured at pH 7.0 (left panel) or supplemented with 1 mM cysteine (right panel).

The human gastrointestinal tract exhibits substantial physicochemical gradients along its length, with pH values ranging from 6.0 to 8.5, oxygen availability varying from microaerobic (pO₂ 2-8 mmHg) to strictly anaerobic, and redox potentials spanning -50 to -350 mV from the duodenum to the rectum.^25–28^ To assess how these environmental parameters influence neurosteroid production, we examined progesterone metabolism by strain DSM 12042 under different pH and redox conditions. Strain DSM 12042 exclusively transformed progesterone into isopregnanolone across all tested pH conditions (4.0-8.0), with maximum production observed at pH 7.0 (**Figure 6B**). We next evaluated the strain’s performance under varying redox conditions: microaerobic atmosphere (5% O₂ in headspace) and anaerobic conditions with or without reducing agents, including cysteine (1 mM, generating a redox potential of approximately -150 mV) and sodium sulfide (1 mM, generating approximately -400 mV).^19^ Isopregnanolone was the sole progesterone-derived product under all conditions tested, although 5α-dihydroprogesterone, the microbial intermediate (**Figure 4A**), appeared under sub-optimal conditions (**Figure S3**). Maximal isopregnanolone production occurred when strain DSM 12042 was cultured in PYG broth (pH 7.0) containing 1 mM cysteine (**Figure 6B**).

### *H. filiformis* strain DSM 12042 modulates neurosteroid profiles in host tissues

We investigated the effects of *H. filiformis* strain DSM 12042 on neurosteroid profiles in different host tissues using C57BL/6 female mice as the model organism. Mice were orally administered progesterone (unlabeled progesterone and [3,4-¹³C₂]progesterone mixed in a 9:1 ratio; 20 mg/kg), strain DSM 12042, and/or high-fat diet. Prior to progesterone treatment, mice received strain DSM 12042 via oral gavage three times within one week (**Figure 7A**). At 90 minutes post-administration of oral progesterone, mice were euthanized and tissues were immediately dissected for steroid profiling by UPLC–HRMS (see **Figure S4** for the detection of ^13^C-labeled neurosteroids in mouse tissues). We quantified bacterial abundance in the mouse cecum by extracting DNA from gastrointestinal contents and performing qPCR (**Figure 7B**). The *baiJ* type 2 gene served as a molecular marker for *Holdemania* because (i) it exhibits restricted distribution among bacteria and is prevalent only in *Holdemania* spp., and (ii) it is associated with 5α-neurosteroid production, a bioprocess rarely observed in microorganisms.

**Figure 7.**
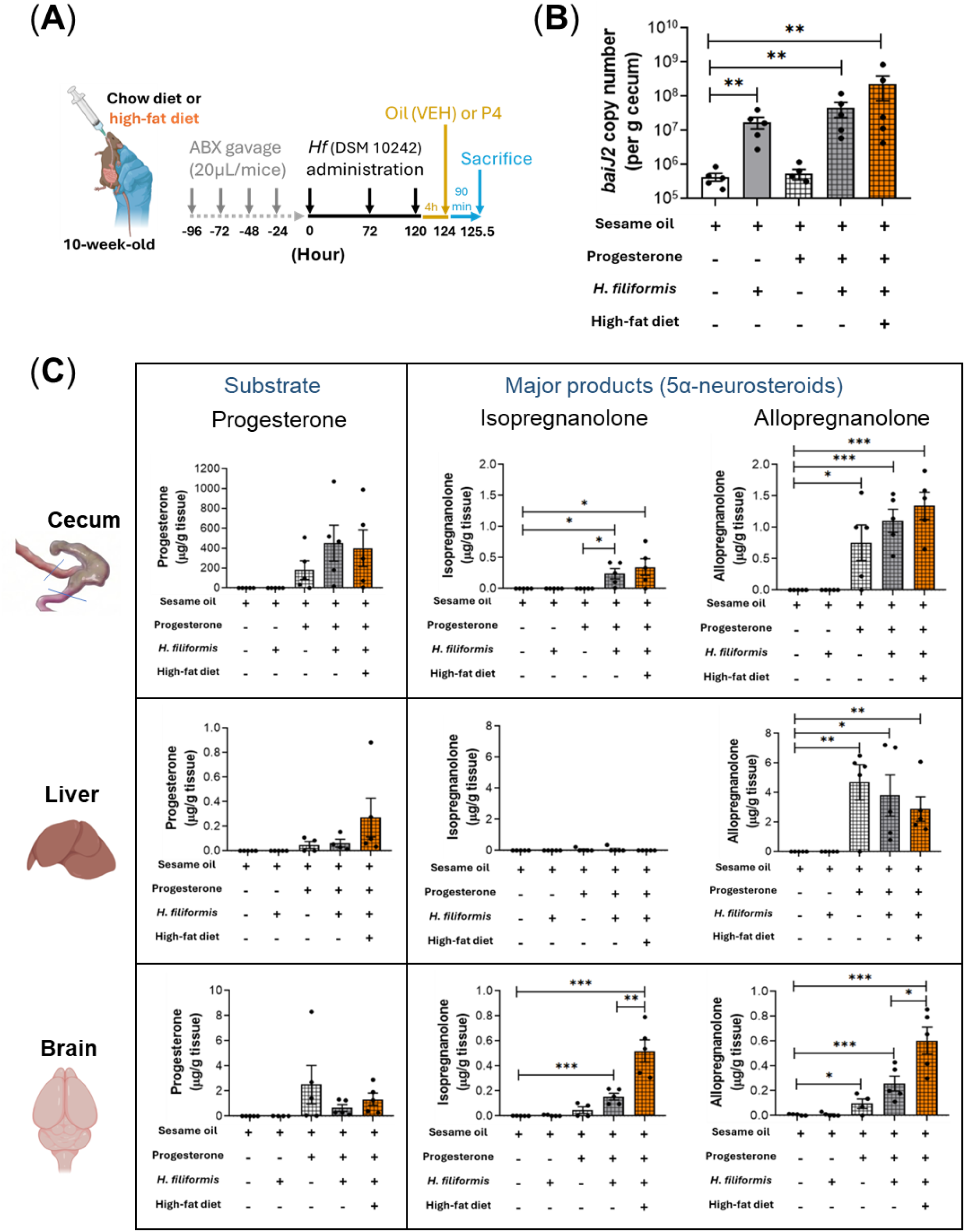
Administration of progesterone (unlabeled progesterone and [3,4-¹³C₂]progesterone mixed in a 9:1 ratio, dissolved in sesame oil) and *H. filiformis* strain DSM 12042 to female mice by oral gavage increases host 5α-neurosteroid levels. (**A**) Experimental workflow for oral gavage administration of strain DSM 12042 and progesterone to female mice. (**B**) Quantification of *H. filiformis baiJ* type 2 gene copy number in mouse cecum using qPCR. Data represent mean ± SEM. Statistical results were calculated with an unpaired nonparametric *t*-test; \*\**p* < 0.01. (**C**) Administration of strain DSM 12042 and progesterone significantly increased isopregnanolone levels in mouse tissues, including cecum and brain. Allopregnanolone was the major neurosteroid detected in mouse liver. Brain 5α-neurosteroid levels, particularly isopregnanolone, were significantly higher in female mice fed a high-fat diet compared with those fed a normal diet. See **Figure S6** for 5β-neurosteroids, including epipregnanolone and pregnanolone. Data represent mean ± SEM. Statistical results were calculated with an unpaired *t*-test; \**p* < 0.05, \*\**p* < 0.01, \*\*\**p* < 0.005.

**Figure 8.**
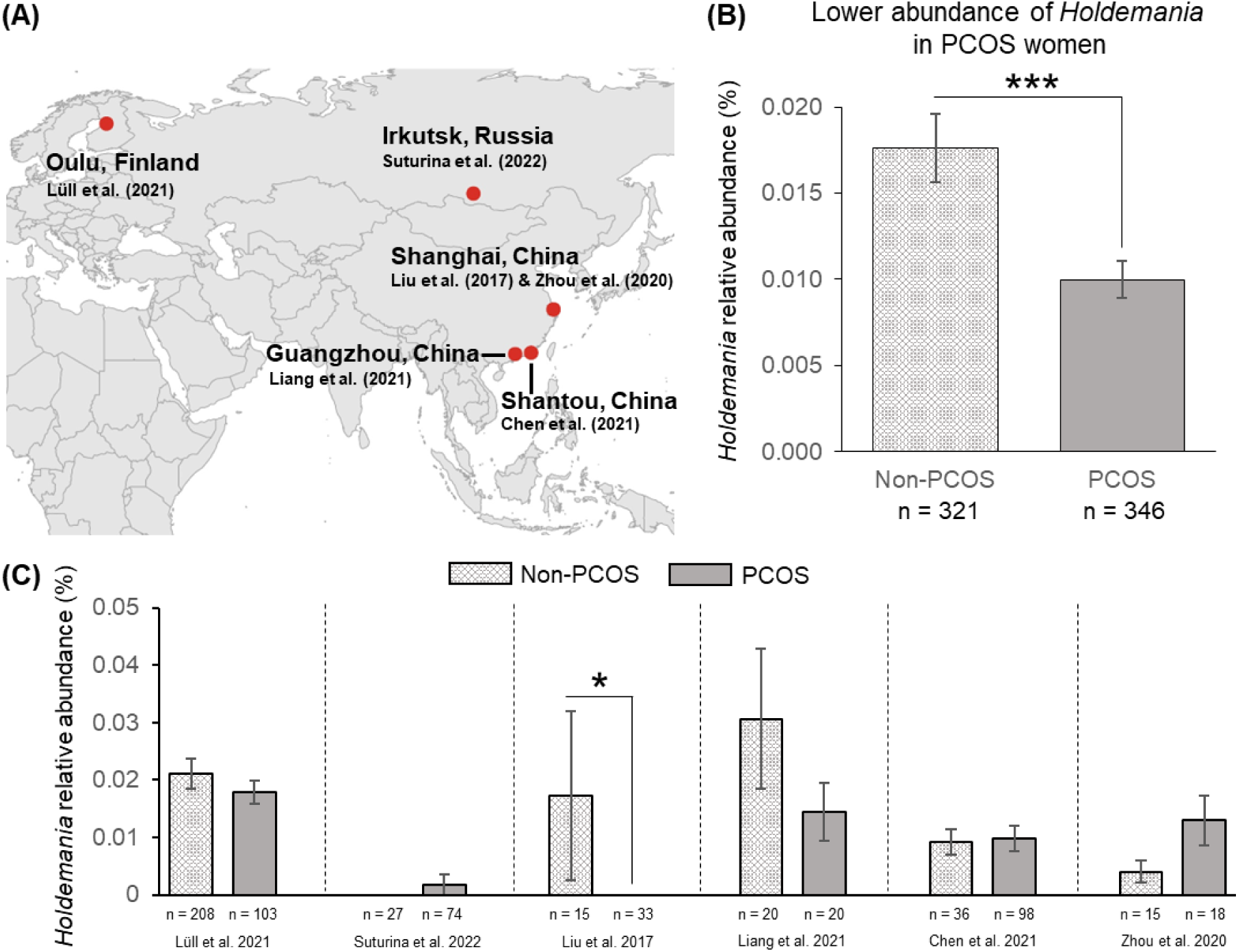
Global meta-analysis reveals reduced *Holdemania* abundance in PCOS patients compared to healthy (non-PCOS) women. Gut microbiota data from non-PCOS women (C; totally 321) and PCOS patients (totally 346) across 6 studies were analyzed. Data represent mean ± SEM. *p < 0.05; ***p < 0.001, Wilcoxon rank-sum test. (**A**) Geographic distribution of the included studies: 1 from Finland, 1 from Russia, and 4 from China. (**B**) *Holdemania* abundance is significantly lower in PCOS patients compared to healthy controls. (**C**) Study-specific *Holdemania* abundance comparisons between healthy controls and PCOS patients.

qPCR analysis using *Holdemania*-specific primers (see **Table S3** for sequences) revealed that cecal *Holdemania* abundance was lowest in mice not administered strain DSM 12042 and was significantly increased in mice receiving this strain (**Figure 7B**). The highest cecal *Holdemania* abundance was observed in mice treated with progesterone, strain DSM 12042, and high-fat diet (up to 1.0 × 10⁹ copies/g tissue). In contrast, total bacterial abundance in the mouse cecum did not differ significantly across treatments (**Figure S5**). These findings indicate that (i) strain DSM 12042 exhibits high survival in the mouse cecum, and (ii) high-fat diet promotes *Holdemania* growth in the host gut.

In the mouse cecum, both isopregnanolone (up to 0.8 μg/g tissue) and allopregnanolone (up to 1.9 μg/g tissue) were the major neurosteroids (**Figure 7C**, upper panel), with levels significantly elevated in mice treated with progesterone and strain DSM 12042. Notably, progesterone treatment alone resulted in a significant increase in cecum allopregnanolone levels but did not appreciably increase those of isopregnanolone. These results suggest the crucial role of *H. filiformis* in gut isopregnanolone production. In mouse liver, allopregnanolone [up to 7.2 μg/g tissue] was the only neurosteroid detected in progesterone-treated mice (**Figure S6**). However, no significant differences in hepatic allopregnanolone levels were observed among mice treated with progesterone alone (4.7 ± 1.2 μg/g tissue), progesterone plus strain DSM 12042 (3.8 ± 1.4 μg/g tissue), or progesterone plus strain DSM 12042 and high-fat diet (2.8 ± 0.8 μg/g tissue) (**Figure 7C**, middle panel). These results indicate the important role of hepatic enzymes in allopregnanolone production. Brain tissue analysis revealed both allopregnanolone [up to 0.9 μg/g tissue] and isopregnanolone [up to 0.8 μg/g tissue] as the major neurosteroids in progesterone-treated mice (**Figure 7C**, bottom panel). Although substantially lower isopregnanolone levels were observed in the mouse cecum, the two 5α-neurosteroids were present in comparable amounts in brain tissue. Their levels were significantly increased following oral treatment with both progesterone and strain DSM 12042. Further increased levels of both 5α-neurosteroids were observed in mice treated with progesterone, strain DSM 12042, and high-fat diet (**Figure 7C**, bottom panel).

### Global meta-analysis of *Holdemania* abundance in PCOS versus non-PCOS women

We analyzed six studies comparing PCOS and non-PCOS groups across geographically diverse populations: Oulu, Finland (PRJNA669650);^29^ Irkutsk, Russia (PRJNA899143);^30^ Shanghai, China (PRJNA341567 and PRJNA634229);^31,32^ Guangzhou, China (PRJNA622999);^33^ and Shantou, China (PRJNA737206).^34^ Meta-analysis revealed significantly lower relative abundance of *Holdemania* in PCOS women (n = 346) compared to non-PCOS women (n = 321) (*p* < 0.001). At the individual study level, Liu et al. (2017)^31^ demonstrated a statistically significant reduction in *Holdemania* abundance in PCOS women (*p* < 0.05). Similarly, Lüll et al. (2021)^29^ and Liang et al. (2021)^33^ both showed trends toward lower *Holdemania* in PCOS women, though these differences did not reach statistical significance.

## DISCUSSION

The gut-brain axis has emerged as a critical bidirectional communication system influencing neurological function, mood regulation, and overall health.^35,36^ Our findings establish that specific gut microbes, particularly *Holdemania* species, serve as substantial producers of 5α-neurosteroids. This discovery fundamentally expands our understanding of neurosteroid biosynthesis beyond exclusively host-mediated pathways and reveals the gut microbiome as a pharmacologically relevant source of neuroactive compounds.

Our metabolomic analyses of progesterone-amended fecal cultures demonstrated that gut microbiota generates the complete spectrum of C₂₁ neurosteroids, with pregnanolone (a 5β-neurosteroid) predominating in most individuals. However, we identified substantial inter-individual variation in neurosteroid production profiles, likely reflecting differences in gut microbial community composition. Notably, allopregnanolone and isopregnanolone—both 5α-neurosteroids with potent GABA_A_ receptor modulating activity^2,3,5^—were detected in several samples, suggesting that certain individuals harbor gut microbes capable of producing therapeutically relevant neurosteroids. Through integrated genomic and metabolomic approaches, we identified *Holdemania* as a key producer of isopregnanolone, mediated by a microbial steroid 5α-reductase (BaiJ homolog) and 3β-hydroxysteroid dehydrogenase/reductase. Phylogenetic analysis revealed that BaiJ-like sequences are predominantly found within Firmicutes (Bacillota), with *Holdemania* species forming a distinct clade (BaiJ type 2) showing 48-50% amino acid identity to the bile salt-degrading *Clostridium scindens* BaiJ. The restricted phylogenetic distribution of steroid 5α-reductase genes thus aligns with our metabolomic findings showing pregnanolone as the dominant microbial metabolite, reflecting the greater prevalence of 5β-reductase pathways in gut bacteria.

The functional characterization of *H. filiformis* strain DSM 12042 demonstrated robust isopregnanolone production across diverse physicochemical conditions mimicking the gastrointestinal environment, including pH ranges of 4.0-8.0 and varying redox potentials. Optimal production occurred at pH 7.0 with mild reducing conditions (1 mM cysteine), corresponding to conditions prevalent in the cecum and colon. This environmental adaptability suggests that *H. filiformis* maintains neurosteroid biosynthetic capacity throughout the intestines. A critical question addressed by our study concerns whether gut-produced neurosteroids reach the brain in pharmacologically relevant concentrations. Using stable isotope tracing with [3,4-¹³C₂]progesterone, we obtained unequivocal evidence for gut-to-brain transport of microbiota-derived neurosteroids. Specifically, ¹³C-labeled isopregnanolone was detected in brain tissue of female mice orally administered labeled progesterone and *H. filiformis*. Notably, the neurosteroid profile in brain tissue more closely resembled that of gut tissues than hepatic tissue, where allopregnanolone was the exclusive neurosteroid detected (**Figure S6**). This finding challenges the traditional model in which hepatically produced allopregnanolone represents the primary source of circulating neurosteroids. Given that *H. filiformis* exclusively produces isopregnanolone from progesterone, the allopregnanolone detected in mouse cecum likely originated from the liver, where hepatic enzymes convert orally administered progesterone to allopregnanolone. This interpretation is further supported by evidence of steroid transport through enterohepatic circulation in rodents, including mice.^18^

Although allopregnanolone levels were significantly higher in the cecum and liver, both 5α-neurosteroids (allopregnanolone and isopregnanolone) were present at similar concentrations in brain tissue. This finding suggests differential transport efficiency, with isopregnanolone exhibiting preferential or more efficient transport through the gut-brain axis compared to its hepatic-derived counterpart, allopregnanolone. The enhanced accumulation of both 5α-neurosteroids in brain tissue of mice receiving high-fat diet warrants particular attention. Dietary lipids may enhance neurosteroid bioavailability through multiple mechanisms, including increased intestinal absorption, altered blood-brain barrier permeability, or modulation of gut microbial composition and metabolism. This observation aligns with emerging evidence linking dietary composition to neurosteroid levels and mood regulation,^37–40^ suggesting that nutritional interventions might potentiate the therapeutic effects of probiotic neurosteroid-producing bacteria.^39,40^

The identification of gut microbiota as alternative neurosteroid sources has profound therapeutic implications. Current neurosteroid-based therapies face significant challenges: allopregnanolone (Zulresso™) requires intravenous administration over 60 hours due to limited oral bioavailability,^41–43^ while chemical synthesis produces racemic mixtures requiring costly purification.^44,45^ At a similar oral administration dose, the colonization efficiency of *H. filiformis* (10⁹ copies/g cecum tissue) is approximately 100-fold higher than that of another steroid-metabolizing gut microbe, *Thauera* sp. strain GDN1 (10⁷ copies/g cecum tissue).^18^ Our findings suggest that colonization with *H. filiformis* or closely related neurosteroid-producing bacteria could provide sustained, endogenous neurosteroid production, potentially offering advantages over conventional pharmacological approaches. Moreover, the substantial inter-individual variation in neurosteroid production profiles observed across our female cohort highlights the importance of personalized approaches to microbiome-based therapeutics. Factors influencing this variation likely include baseline gut microbial composition, dietary habits, antibiotic exposure history, host genetics affecting gut physiology, and hormonal status. Understanding these determinants will be essential for identifying individuals most likely to benefit from probiotic interventions with neurosteroid-producing bacteria.

Patients with polycystic ovary syndrome (PCOS) experience anovulation or chronic anovulation, resulting in absent or infrequent progesterone production and reduced progesterone exposure.^17,46^ They also frequently exhibit alterations in gut microbiota composition and a high prevalence of mood disorders.^17,47^ Our current data reveal the progesterone-dependent growth of *Holdemania* species, particularly *H. filiformis*. Given this relationship, the significant decrease in *Holdemania* abundance observed in the PCOS population is not surprising. The reduced abundance of *Holdemania* in certain disease states, such as PCOS, as demonstrated by our comparative microbiome analyses, raises the possibility that depletion of neurosteroid-producing bacteria contributes to disease pathogenesis. Future studies should investigate whether *Holdemania* abundance correlates with mood symptoms, circulating neurosteroid metabolite levels, or treatment response to conventional antidepressant therapies. Such investigations could identify microbiome-based biomarkers for stratifying patients and personalizing therapeutic interventions.

## CONCLUSION

This study establishes the gut microbiome as a significant source of 5α-neurosteroids with therapeutic potential for neurological and psychiatric disorders. We identified *Holdemania* species as a major producer of isopregnanolone through microbial steroid 5α-reductase and 3β-hydroxysteroid dehydrogenase/reductase activities. Using stable isotope tracing, we demonstrated gut-to-brain transport of microbiota-derived neurosteroids, challenging the traditional paradigm that hepatic synthesis represents the exclusive source of circulating neurosteroids.^48^ The distinct neurosteroid profiles observed in liver (allopregnanolone predominant) versus gut and brain tissues (both allopregnanolone and isopregnanolone predominant in *H. filiformis*-treated mice) suggest that gut microbiota contribute substantially to central nervous system neurosteroid pools. The enhancement of brain 5α-neurosteroid levels by high-fat diet further indicates that nutritional interventions may potentiate microbial neurosteroid production and bioavailability. Our phylogenetic analyses revealed that steroid 5α-reductase genes are restricted to specific bacterial lineages, predominantly within Firmicutes, explaining the inter-individual variation in 5α-neurosteroid production capacity. The identification of *Holdemania* species harboring both 5α-reductase and 3β-hydroxysteroid dehydrogenase genes positions this genus as a particularly promising target for probiotic development.

These findings have immediate translational implications. *H. filiformis* represents a promising probiotic candidate for enhancing endogenous neurosteroid production through gut-brain axis modulation, potentially offering advantages over current neurosteroid-based therapies including improved bioavailability, sustained production, and oral administration. Future clinical studies should evaluate the safety, tolerability, and efficacy of *H. filiformis* supplementation in patients with mood disorders, postpartum depression, or other conditions associated with neurosteroid deficiency. Beyond immediate therapeutic applications, our work fundamentally expands understanding of the gut-brain axis by revealing that gut microbiota serve as endocrine organs capable of producing neuroactive steroids that reach the brain in pharmacologically relevant concentrations. This discovery opens new avenues for investigating microbiome contributions to neurological health and disease, and for developing novel microbiome-based interventions for neuropsychiatric conditions. The integration of microbial genomics, metabolomics, and in vivo functional studies exemplified in this work provides a roadmap for discovering and characterizing additional microbial contributions to host physiology and for translating these discoveries into clinical applications.

## MATERIALS AND METHODS

### Chemicals and bacterial strains

[3,4-¹³C₂]progesterone (98%) was purchased from Sigma-Aldrich (St. Louis, MO, USA). Four neurosteroids were purchased from Steraloids (Newport, RI, USA): 3α-hydroxy-5α-pregnan-20-one (allopregnanolone), 3β-hydroxy-5α-pregnan-20-one (isopregnanolone), 3α-hydroxy-5β-pregnan-20-one (pregnanolone), and 3β-hydroxy-5β-pregnan-20-one (epipregnanolone). All other chemicals were of analytical grade and sourced from Mallinckrodt Baker (Phillipsburg, NJ, USA), Merck Millipore (Burlington, VT, USA), and Sigma-Aldrich. Strains *H. filiformis* DSM 12042 and *H. massiliensis* AP2 were acquired from the Leibniz Institute DSMZ (Germany).

### Participants, protocols, and sample collection

Human fecal sample collection was performed as described in a previous study,^17^ with approval from the local ethics committee (Clinical Trial/Research Approval NTUH-REC no.:202103046RINB). This study included infertile female patients undergoing hormonal endometrial preparation before frozen-thawed embryo transfer. Hormone replacement therapy began on menstrual cycle day 3. Estradiol valerate was administered in increasing doses: 4 mg/day (days 3-8), 8 mg/day (days 9-11), and 12 mg/day (from day 12) until optimal endometrial thickness was achieved. Two days before embryo transfer, transvaginal progesterone gel (8% Crinone; Merck) was administered at 90 mg/day for 2 days, then increased to 180 mg/day for 14 days. Oral progesterone (Utrogestan, 600 mg/day) was initiated on embryo transfer day. Participants provided fecal samples one day before starting progesterone treatment.

### Identification of microbial progestogenic metabolites in fecal cultures

Fresh fecal samples were collected from infertile women aged 32–43 years (see **Table S1**) who underwent oral progesterone therapy for endometrial preparation and thawed embryo transfer. Samples (approximately 0.5 g each) were anaerobically incubated with progesterone (1 mM) in either chemically defined medium containing mineral salts (DCB-1, 100 mL) or Brain Heart Infusion (BHI) medium (100 mL) at 37°C in darkness. Subsequently, 17α-ethinylestradiol (final concentration: 50 μM), which is not metabolized by gut microbiota, was added as internal control. Daily samples were extracted from progesterone-enriched fecal cultures, and progesterone-derived microbial products were isolated through dual extraction with ethyl acetate. After complete solvent evaporation, residues were reconstituted in 50 μL methanol, and progestogenic metabolites were identified using ultraperformance liquid chromatography–atmospheric pressure chemical ionization–high-resolution mass spectrometry (UPLC–APCI–HRMS). To determine temporal changes in bacterial community structures within fecal cultures, bacterial cells were collected by centrifugation (10,000 × g for 10 min at 4°C). Bacterial DNA was extracted using a QIAamp PowerFecal Pro DNA kit (Qiagen, Hilden, Germany). DNA concentration was determined using either a NanoDrop ND-1000 spectrophotometer or Qubit dsDNA Assay kit (Invitrogen, Thermo Fisher Scientific, Waltham, MA, USA). Bacterial 16S rRNA was amplified by PCR, and amplicons were sequenced using the PacBio platform.

### UPLC–APCI–HRMS

Progesterone and its derivatives were analyzed using UPLC–HRMS, consisting of a UPLC system coupled to an APCI-MS system. Metabolite separation was performed on a reversed-phase C18 column (ACQUITY UPLC BEH C18; 1.7 μm, 100 × 2.1 mm; Waters, Milford, MA, USA) at 0.45 mL/min with column oven at 50°C. The mobile phase comprised solution A (0.1% formic acid (v/v) in 2% acetonitrile (v/v)) and solution B (0.1% formic acid (v/v) in acetonitrile (v/v)). Gradient elution increased solution B from 40% to 55% over 9 minutes, then to 100% within 1 minute. Mass spectrometric data were collected in positive ionization mode with parent scan range of 100-500 *m/z*. Capillary and APCI vaporizer temperatures were 120°C and 395°C, respectively. Sheath, auxiliary, and sweep gas flow rates were 40, 5, and 2 arbitrary units. Source voltage was 6 kV with 15 μA current. Individual adduct ion compositions were determined using Xcalibur Software (Thermo Fisher Scientific).

### Thin-layer Chromatography (TLC)

Steroid standards and products were separated on silica gel-coated aluminum TLC plates (Silica gel 60 F254, 0.2 mm thickness, 20 × 20 cm; Merck) using dichloromethane: ethyl acetate: ethanol (14:4:0.05, v/v/v) as the mobile phase. Steroids were visualized under UV light at 254 nm or by spraying the TLC plates with 30% (v/v) H₂SO₄ followed by heating at 100°C for 1 min in an oven. In some cases, spots corresponding to individual steroids were quantified using ImageJ software (https://imagej.net/ij/).

### Routine cultivation and physiological characterization of *H. filiformis*

*H. filiformis* strain DSM 12042 was routinely cultured in PYG broth with or without progesterone (1 mM, terminal electron acceptor) under strictly anaerobic conditions. To determine optimal physicochemical conditions for progesterone-to-isopregnanolone conversion, strain DSM 12042 was cultured in PYG broth supplemented with 1 mM progesterone under varying pH (4.0–8.0 in 1.0-unit increments) or redox conditions: microaerobic (5% O₂), anaerobic (0% O₂), anaerobic with the mild reducing agent L-cysteine (1 mM, approximately −150 mV), or anaerobic with the strong reducing agent Na₂S (1 mM, approximately −400 mV). Steroids were extracted from bacterial cultures with ethyl acetate and quantified by UPLC–HRMS.

### *Holdemania*-mediated progesterone-to-isopregnanolone conversion

To compare progesterone-to-isopregnanolone conversion by *H. filiformis* DSM 12042 and *H. massiliensis* AP2, pre-cultures were prepared by inoculating 10 mL of PYG medium with a 2% (v/v) inoculum under strict anaerobic conditions. Following overnight incubation at 37°C, cultures were diluted to an OD₆₀₀ of approximately 0.3 using fresh PYG medium. One milliliter of each normalized culture was then used to inoculate 100 mL of PYG medium supplemented with 1 mM progesterone. Cultures were incubated at 37°C with agitation at 220 rpm for 24 h under anaerobic conditions. Culture samples (1 mL) were collected in triplicate at 24 h and immediately processed for metabolite extraction. Each sample was extracted twice with 0.8 mL of ethyl acetate, with vortexing for 60 s and subsequent centrifugation (6,000 ×*g*, 2 min) to facilitate phase separation. The combined organic phases were transferred to clean tubes and dried under vacuum. Dried residues were reconstituted in 100 µL of methanol and analyzed by UPLC–HRMS.

### Molecular biological methods

Bacterial genomic DNA was extracted with Presto Mini gDNA Bacteria Kit (Geneaid, New Taipei City, Taiwan). Full-length 16S rRNA genes were amplified using universal primers: Forward (27F): 5’-AGAGTTTGATCMTGGCTCAG-3’; Reverse (1492R): 5’- TACGGYTACCTTGTTACGACTT-3’. PCR amplification used 25-μL reactions containing nuclease-free H_2_O, Invitrogen Platinum Hot Start PCR 2X Master Mix (Thermo Fisher Scientific), 200 nM of each primer, and 10-30 ng template DNA. PCR products were verified by 1.5% TAE-agarose gel electrophoresis with SYBR Green I nucleic acid stain (Invitrogen, Thermo Fisher Scientific) and purified using GenepHlow Gel/PCR Kit (Geneaid).

### PacBio sequencing of bacterial 16S rRNA gene amplicons

Bacterial genomic DNA was extracted using a QIAamp PowerFecal DNA Kit (Qiagen) and quantified with a Qubit 4.0 Fluorometer (Thermo Scientific). The full-length 16S rRNA gene (V1-V9 regions) was amplified using barcoded universal primers. Each primer contained a 5′-phosphorylated buffer sequence (GCATC), a 16-base barcode, and degenerate 16S-specific sequences: Forward: 5′Phos/GCATC-[16-base barcode]-AGRGTTYGATYMTGGCTCAG-3; Reverse: 5′Phos/GCATC-[16-base barcode]-RGYTACCTTGTTACGACTT-3′; (where M = A/C; R = A/G; Y = C/T). PCR amplification was performed using KAPA HiFi HotStart ReadyMix (Roche) with 2 ng template DNA. Cycling conditions were: 95°C for 3 min; 20-30 cycles of 95°C for 30 s, 57°C for 30 s, and 72°C for 60 s; followed by 72°C for 5 min. PCR products were verified by 1% agarose gel electrophoresis. Samples showing distinct ∼1,500 bp bands were purified using AMPure PB Beads for PacBio library preparation. Raw PacBio fastq files were deposited in NCBI SRA: DCB samples under BioProject PRJNA1088978 (accession numbers SRR30505914-SRR30505967) and BHI samples under BioProject PRJNA1196623 (accession numbers SRR31672780-SRR31672827).

### PacBio read quality filtering and taxonomic classification

Raw PacBio reads were processed using QIIME2-DADA2 pipeline (version 2023.5).^49,50^ Sequences shorter than 1,000 bp or longer than 1,600 bp, including primers and chimeric sequences, were removed. ASVs were classified using bacterial 16S rRNA RefSeq sequences from NCBI nucleotide database. Classification thresholds: species (≥98.7%), genus (≥94.5%), family (≥86.5%), order (≥82.0%), class (≥78.5%), and phylum (≥75.0%).^51–53^ Samples were rarefied to 10,075 reads and ASV counts converted to percentages.

### Bacterial community structure analysis of progesterone-treated gut microbiota

Fecal samples from 14 female patients were anaerobically cultured in BHI or DCB-1 broth with 1 mM progesterone. Bacterial community structure differences were analyzed using NMDS. Differential abundance analysis between control (without progesterone) and progesterone-supplemented samples was performed using the Wilcoxon rank-sum test at the phylum level and DESeq2 at the genus level.^54^

### Identification and comparative analysis of BaiJ-like sequences

The BaiJ from *Clostridium scindens* VPI 12708 (WP_025644103.1) was used as a query in BLAST searches against Bacillota genomes in the NCBI database (ClusteredNR setting). Bacterial strains with deduced amino acid sequences showing >40% identity to BaiJ were identified, and their genomes were downloaded. These proteins were compared via BLAST to established hydroxysteroid dehydrogenase/reductase (HSDR) references: BaiA1/3 and BaiA2 from *C. scindens* DSM 5676 (3α-HSDR; Meibom et al., 2024),^21^ Rumgna_02133 (A7B3K3.1) and Rumgna_00694 (A7AZH2.1) from *Ruminococcus gnavus* DSM 114966 (3α- and 3β-HSDR, respectively), and ci2349 from *C. innocuum* DSM 24813 (WP_100933575.1; 3β-HSDR).^55^ Results were visualized in a heatmap displaying sequence identity between bacterial proteins and references. Proteins were arranged in descending order of identity to BaiJ (>40% threshold). A blue-white-red gradient indicates sequence identity percentage, with sequences showing >90% coverage displayed in bold. Note: *C. innocuum* 6_1_30 (GCA_000183585.2) corresponds to *Clostridium* sp. HGF2 in the NCBI genome database.

### Phylogenetic tree of BaiJ protein sequences across major gut bacterial phyla

Sequence retrieval and selection for phylogenetic analysis. The *C. scindens* BaiJ protein sequence was used as a query for BLAST searches against four major bacterial phyla (Actinomycetota, Bacillota, Bacteroidota, and Pseudomonadota) in the NCBI database using the ClusteredNR setting. The top 100 protein sequences from each phylum were downloaded and subjected to quality filtering. Sequences shorter than 400 amino acids were excluded to ensure analytical robustness. Multiple sequence alignment and filtering. Multiple sequence alignment was performed using MAFFT, followed by quality filtering with trimAl to remove low-quality sequences and poorly aligned regions.^56,57^ Sequences with fewer than 50% good positions were excluded (−seqoverlap 50), along with alignment columns containing less than 50% residue overlap (−resoverlap 0.5) or more than 95% gaps (−gt 0.05). This filtering process resulted in 390 high-quality aligned sequences for phylogenetic analysis.

Phylogenetic reconstruction was performed using IQ-TREE version 3.0.1 with the LG + I + G4 + F evolutionary model to account for rate heterogeneity across sites.^58^ Tree reliability was assessed using 1,000 ultrafast bootstrap replicates and 1,000 replicates of the SH-aLRT (Shimodaira-Hasegawa approximate Likelihood Ratio Test). Trees were visualized using the Interactive Tree of Life (iTOL) platform.^59^

### Quantification of baiJ and 16S rRNA in cecal content by quantitative PCR (qPCR)

The copy numbers of *baiJ* and bacterial 16S rRNA were determined using quantitative PCR (qPCR). Calibration curves were obtained through 10-fold serial dilution of the full-length PCR products of *baiJ* from *Holdemania filiformis* and universal bacterial 16S rRNA. Primer sequences are provided in **Table S3**.

### *Holdemania filiformis* administration in mice

C57BL/6J female mice aged 10 weeks were obtained from the Animal Center of the College of Medicine of National Taiwan University (Taipei, Taiwan) and maintained under standard conditions according to health guidelines for the care and use of experimental animals. All experiments were approved by the local ethics committee (IACUC No. 20220423). Before *H. filiformis* administration, the mice were fed with a standard chow diet or a high-fat diet (60% high-calorie diet, ResearchDiet, USA) and treated with ABX solution (containing amphotericin-B 0.1 mg/mL, ampicillin 10 mg/mL, neomycin 10 mg/mL, metronidazole 10 mg/mL, and vancomycin 5 mg/mL) for 4 days through oral gavage to deplete the original microbiota. Mice fed with a normal chow diet and with similar body weight (18–20 g) were randomly assigned to treatment groups. For the administration of *H. filiformis* strain DSM 12042, approximately 1 × 10^9^ CFUs [suspended in 100 uL of phosphate-buffered saline (PBS)] were fed to each mouse through oral gavage at day 0, 3, and 5. Progesterone was prepared as a mixture of unlabeled progesterone and [3,4-¹³C₂]progesterone at a 9:1 ratio and dissolved in sesame oil (Sigma-Aldrich). Mice were administered a progesterone mixture (with a dose of 20mg/kg) after 4 hours of the final *H. filiformis* administration by oral gavage. Subsequently, the mice were euthanized after 90 minutes of progesterone administration (anesthetized with 3% isoflurane). Tissue samples, including cecum, liver, and brain, were immediately collected and stored at −80°C before steroid profile analysis.

### Tissue steroid extraction

Tissue samples from the brain, cecum, and liver were collected into pre-weighed, low-protein binding microcentrifuge tubes (Protein LoBind, Eppendorf, Germany). Following 48-h lyophilization (KINGMECH FD3-12P, Taiwan), the dried tissues were pulverized using disposable pestles to minimize transfer loss. Each sample was spiked with 19-nortestosterone as an internal standard and rehydrated with 500 µL of ultrapure water. Neurosteroids were extracted twice with 900 µL of ethyl acetate through 10-min vortexing and centrifugation (16,000 ×*g*, 5 min). The combined organic phases were evaporated to dryness under vacuum and reconstituted in 300 µL of LC-MS grade methanol. To reduce matrix interference from lipid-rich tissues, samples were incubated at -20°C for 2 h to precipitate lipids, followed by centrifugation (16,000 ×*g*, 10 min, 4°C). The supernatant was transferred to autosampler vials with glass inserts for UPLC-HRMS analysis. Analytical consistency was monitored by internal standard recovery. Statistical analyses and data visualization were performed using GraphPad Prism version 10 (GraphPad Software, San Diego, CA, USA).

### Comparative meta-analysis

Studies were selected based on three criteria: (1) publicly available datasets in NCBI, (2) properly labeled non-PCOS and PCOS groups in the metadata, and (3) 16S rRNA sequencing data rather than shotgun metagenomic data. Six studies met these criteria: Oulu, Finland (PRJNA669650, Lüll et al., 2021);^29^ Irkutsk, Russia (PRJNA899143, Suturina et al., 2022);^30^ Shanghai, China (PRJNA341567, Liu et al., 2017; PRJNA634229, Zhou et al., 2020);^31,32^ Guangzhou, China (PRJNA622999, Liang et al., 2021);^33^ and Shantou, China (PRJNA737206, Chen et al., 2021).^34^ Most datasets targeted the V3-V4 region of the 16S rRNA gene, except for the Russian cohort (V1-V3) and one Chinese cohort (V4, Liang et al., 2021).^33^ In total, 667 samples were analyzed [non-PCOS (n=321) and PCOS (n=346)]. Fastq files from each study were quality-filtered and processed independently using the QIIME2-DADA2 pipeline. Count data were then combined and normalized by proportional abundance rather than rarefaction to preserve low-abundance taxa such as *Holdemania*. Differential abundance analysis was performed for *Holdemania*.

### Statistical analyses

All experiments were performed at least in triplicate. Data normality was assessed using the Shapiro-Wilk test. Normal distributions were analyzed using Welch’s t-test; non-normal distributions using Wilcoxon rank sum test. Significance was set at p < 0.05. Bacterial community structure changes were assessed by PERMANOVA (adonis) using R package vegan. Differential abundance analysis was performed using DESeq2 at the genus level.^54^

## Acknowledgements

We thank the Small Molecule Metabolomics core facility sponsored by the Institute of Plant and Microbial Biology, Academia Sinica (Taiwan), for its UPLC–HRMS analysis (AS-CFII-111-218).

## Author contributions

G.-J.B.M., M.-J.C., T.-H.H., and Y.-R.C. designed the research; G.-J.B.M., C.-H. C., T.-Y.W., R.G.G., C.-T.H., G.-L.W., Y.-C.L., Y.-A.T., Y.-Y.C., Y.-L.L, P.-T.L., and Y.-L.T. conducted the research; M.-J.C., and T.-H.H. contributed new reagents and analytic tools; G.-J.B.M., C.-H. C., T.-Y.W., M.-J.C., T.-H.H., and Y.-R.C. analyzed the data; and G.-J.B.M., M.-J.C., T.-H.H., and Y.-R.C. composed the manuscript.

## Data availability

Raw PacBio fastq files were deposited in NCBI SRA under two BioProjects: PRJNA1088978 (accession numbers SRR30505914-SRR30505967) for bacterial 16S rRNA amplicons extracted from DCB-1 cultures, and PRJNA1196623 (accession numbers SRR31672780-SRR31672827) for the BHI cultures. Previously published datasets used for comparative meta-analysis were obtained from the following BioProjects: PRJNA669650 (Lüll et al., 2021), PRJNA899143 (Suturina et al., 2022), PRJNA341567 (Liu et al., 2017), PRJNA634229 (Zhou et al., 2020), PRJNA622999 (Liang et al., 2021), and PRJNA737206 (Chen et al., 2021). All data are available in the main text or the supplementary materials.

## Ethics approval

Human fecal sample collection was approved by the Clinical Trial /Research Approval NTUH-REC No.:202103046RINB. This animal study conducted in accordance with the requirements of the Taiwan Animal Protection Law on Scientific Application of Animals and the Institutional Animal Care and Use Committee (IACUC) of National Taiwan University College of Medicine (IACUC No. 20220423).

## Supplementary Information

### Additional file 1

#### SI Figures

**Fig. S1.** Bacterial phyla significantly enriched by progesterone treatment. Bars represent mean relative abundance ± SEM (n = 14). (−) control without progesterone; (+) progesterone-supplemented. Statistical significance was determined by Wilcoxon rank-sum test: \**p* < 0.05, \*\**p* < 0.01, \*\*\**p* < 0.001.

**Fig. S2.** Heatmap showing the top 30 bacterial genera significantly enriched in progesterone-treated fecal cultures compared to untreated controls (*p* < 0.05). Differential abundance was calculated using DESeq2, and normalized abundance values are displayed as Z-scores. Genera are arranged vertically in ascending order by p-value, with the most significantly enriched genus at the top (rank 1).

**Figure S3.** Metabolic conversion of progesterone by *H. filiformis* strain DSM 12042. Thin-layer chromatography (TLC) identifies isopregnanolone as the primary neurosteroid derivative of progesterone. The microbial intermediate, 5α-dihydroprogesterone (DHP), is detectable under specific growth conditions. Cultures were grown in PYG broth supplemented with 1 mM progesterone. Optimal conversion to isopregnanolone occurred at pH 7.0 in the presence of 1 mM cysteine.

**Figure S4.** Identification of ^13^C-labled 5α-neurosteroids in the mouse brain.The C57BL/6 female mice were orally administered progesterone (unlabeled progesterone and [3,4-¹³C₂]progesterone mixed in a 9:1 ratio; 20 mg/kg), strain DSM 12042, and high-fat diet. Prior to progesterone treatment, mice received strain DSM 12042 via oral gavage three times within one week. At 90 min after oral progesterone administration, mice were euthanized and steroid profiles in tissue samples were determined by UPLC–HRMS.

**Figure S5.** Total bacterial 16S rRNA gene copy number in mouse cecum did not differ significantly across treatments. The 16S rRNA gene was quantified by qPCR using universal primers (see **Table S3** for sequences). Data represent mean ± SEM. Statistical results were calculated with unpaired nonparametric *t*-test.

**Figure S6.** Profiling of C21 neurosteroids in mouse tissues across treatments. Data are represented as mean ± SEM. Abbreviations: Iso Preg, isopregnanolone; Allo Preg, allopregnanolone; Preg, pregnanolone; Epi Preg, epipregnanolone.

#### SI Tables

**Table S1.** Physiological characteristics of 14 women with infertility (with progesterone administration).

**Table S2.** UPLC-HRMS patterns of individual progesterone and four major neurosteroids.

**Table S3.** Oligonucleotides used in this study.

### SI Appendix

**Appendix S1.** DNA sequences of bacterial genes involved in steroid 5α-reduction.

